# ADENYLATE CYCLASE 3 MEDIATES CAROTID BODY ACTIVATION AND AUTONOMIC DYSFUNCTION IN A SLEEP APNEA MODEL

**DOI:** 10.1101/2024.09.24.614747

**Authors:** Ying-Jie Peng, Jayasri Nanduri, Ning Wang, Xiaoyu Su, Matthew Hildreth, Nanduri R. Prabhakar

## Abstract

Patients with obstructive sleep apnea (OSA) experience chronic intermittent hypoxia (CIH). OSA patients and CIH-treated rodents exhibit autonomic dysfunction, characterized by overactive sympathetic nervous system and hypertension, mediated through hyperactive carotid body (CB) chemoreflex. Activation of olfactory receptor 78 (Olfr78) by hydrogen sulfide (H_2_S) is implicated in CB activation and autonomic responses to CIH, but the downstream signaling pathways remain unknown. Given that odorant receptor signaling is coupled to adenylyl cyclase 3 (Adcy3), we hypothesized that Adcy3-dependent cAMP contributes to CB and autonomic responses to CIH. Our findings show that CIH increases cAMP levels in the CB, a response absent in *Adcy3*, *Cth*, and *Olfr78* null mice. CBs from *Cth* and *Olfr78* mutant mice lacked persulfidation response to CIH, indicating that Adcy3 activation by CIH requires Olfr78 activation by H_2_S. CIH also enhanced glomus cell Ca^2+^ influx, an effect absent in *Cnga2* and *Adcy3* mutants, suggesting that CIH-induced cAMP mediates enhanced Ca^2+^ responses through cyclic nucleotide-gated channels. Furthermore, *Adcy3* null mice did not exhibit neither CB activation nor autonomic dysfunction by CIH. These results demonstrate that Adcy3-dependent cAMP is a downstream signaling pathway to H_2_S/Olfr78, mediating CIH-induced CB activation and autonomic dysfunction.

---

Patients with obstructive sleep apnea (OSA) experience chronic intermittent hypoxia (CIH) due to periodic disruptions of breathing during sleep. OSA patients are prone to autonomic dysfunction characterized by elevated sympathetic nerve activity (SNA) and hypertension (Somers *et al*., 1995; Nieto *et al*., 2000; Peppard *et al*., 2000; Dempsey *et al*., 2010). Clinical studies suggest that CIH experienced during OSA during sleep (i.e., hypoxic burden) is a key predictor of disease outcomes of OSA (Redline *et al*., 2023).

Carotid bodies (CB) are the sensory organs for monitoring blood oxygen levels. Hypoxemia (reduced blood oxygen) activates CB neural activity triggering chemoreflex, which is a potent regulator of SNA and BP. Autonomic dysfunction caused by OSA is attributed to an enhanced CB chemoreflex, as indicated by a) heightened CB chemoreflex in OSA patients (Hedner *et al*., 1992), and b) the absence of autonomic changes in OSA patients who have undergone CB resection (Somers & Abboud, 1993). Rodents subjected to CIH, modeled after blood oxygen profiles in obstructive sleep apnea (OSA), exhibit enhanced CB sensitivity to hypoxia and prolonged activation of the CB following acute intermittent hypoxia (simulating apneic episodes) known as sensory long-term facilitation (sLTF). The sLTF may contribute to daytime activation of SNA in OSA patients (Prabhakar, 2013). Understanding the mechanisms underlying CB activation by CIH could potentially lead to novel therapeutic interventions to reduce CB hyperactivity thereby alleviating autonomic dysfunction caused by OSA.

Glomus cells, the primary O₂-sensing cells of the carotid body (CB), express cystathionine γ-lyase (CSE), a H_2_S synthesizing enzyme (Peng *et al*., 2010), and the gene encoding olfactory receptor 78 (Olfr78), a G-protein-coupled receptor (Chang *et al*., 2015; Zhou *et al*., 2016). CIH treated rodents treated exhibit elevated H₂S levels in the CB(Yuan *et al*., 2016). H₂S activates Olfr78 in the CB through persulfidation of Cys ^240^ (Peng *et al*., 2023). Mutant mice lacking genes encoding either *Cth* (encoding CSE) or *Olfr78* show an absence of CB activation (Peng *et al*., 2010; Yuan *et al*., 2015; Peng *et al*., 2020; Peng *et al*., 2023). While these findings suggest that Olfr78 activation by H₂S mediates CB activation by CIH, the signaling downstream of the H₂S- Olfr78 pathway remains unknown.

Olfr78 belongs to the family of odorant receptors (OR). Signaling by ORs involves the generation of cyclic adenosine monophosphate (cAMP) by adenyl cyclase-3 (Adcy3) encoded by the gene *Adcy3* and the activation of the cyclic-nucleotide-gated channel Cnga2 by cAMP. Glomus cells express Adcy3 and Cnga2 proteins (Peng *et al*., 2023). We hypothesized that CIH-evoked CB activation requires increased cAMP generation by Adcy3 and activation of Cnga2, leading to CB hyperactivity and activation of SNA and hypertension. We tested this possibility in CIH- treated mice with targeted deletion of *Adcy3* in tyrosine hydroxylase (TH)-positive CB glomus cells and in mice with heterozygous deficiency of *Cnga2*. Our results showed that CIH increases cAMP levels in the glomus tissue, CB activation, increased plasma catecholamines (an index of sympathetic activation), and elevated blood pressure in wild-type mice. We further found that CIH augments Ca^2+^ influx in wild-type glomus cells but not in CIH-treated *Adcy3* or *Cnga2* mutant mice. Adcy3 null mice showed remarkable absence of CIH-induced CB activation, elevated plasma catecholamines and hypertension.

## RESULTS

### cAMP response of carotid body (CB) to CIH

*Olf78* null mice exhibit absence of CB activation by CIH (Peng *et al*., 2021). Given that olfactory receptor signaling is coupled to Adcy3, we hypothesized that CIH increases cAMP through activation of Adcy3. This possibility was assessed by measuring cAMP levels by ELISA assay in CBs of CIH treated wild-type and in mice with targeted disruption of *Adcy3* in TH positive glomus cells of the carotid body (*Adcy3*^-/-^). Mice treated with room air served as controls. CIH increased cAMP abundance in CBs of wild type mice (Fig.1). In contrast, cAMP response to CIH was absent in *Adcy3* mutant CBs (Fig. 1).

**Figure 1.**
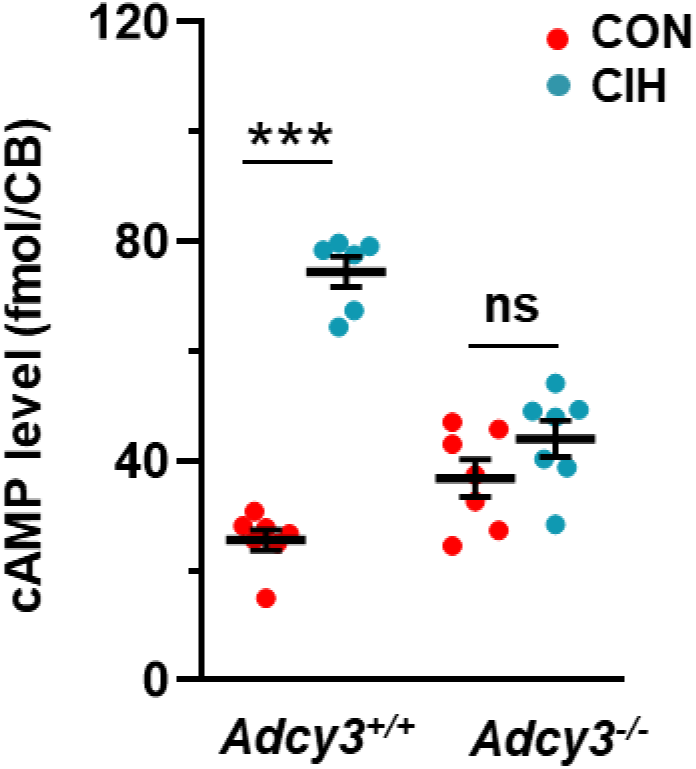
Chronic intermittent hypoxia (CIH) increases Adcy3-dependent cAMP in carotid bodies. cAMP abundance was measured in carotid bodies from mice of the indicated genotypes after CIH treatment for 10d or room air (CON). Data are presented are individual data along with mean ± SEM (2 carotid bodies were pooled from one mouse for each experiment). *** *P* < 0. 001; ns, *P* > 0.05; two-way ANOVA followed by Holm-Sidak test.

We next investigated how CIH activates Adcy3. CIH had no effect on *Adcy3* mRNA levels in wild-type mice (Fig. 1-figure supplement 1). CSE-generated H_2_S activates Olfr78 through persulfidation at the Cys 240 residue (Peng *et al*., 2023). We hypothesized that if CIH increases H_2_S in the CB, it should result in increased persulfidation of Olfr78, and this response should be absent in *Cth* (encoding CSE) and *Olfr78* null mice. To test this possibility, we monitored persulfidation using histochemistry of CB sections and measured cAMP levels in the CBs of wild-type, *Cth*, and *Olfr78* mutant mice. Our results showed that CIH increased persulfidation in wild-type CBs, as indicated by an increased Cy5 signal. This response was absent in *Cth* and *Olfr78* mutant CBs (Fig. 2A). Additionally, CIH increased cAMP levels in wild-type CBs, but not in the CBs of *Cth* and *Olfr78* mutants (Fig. 2B-C).

**Figure 2.**
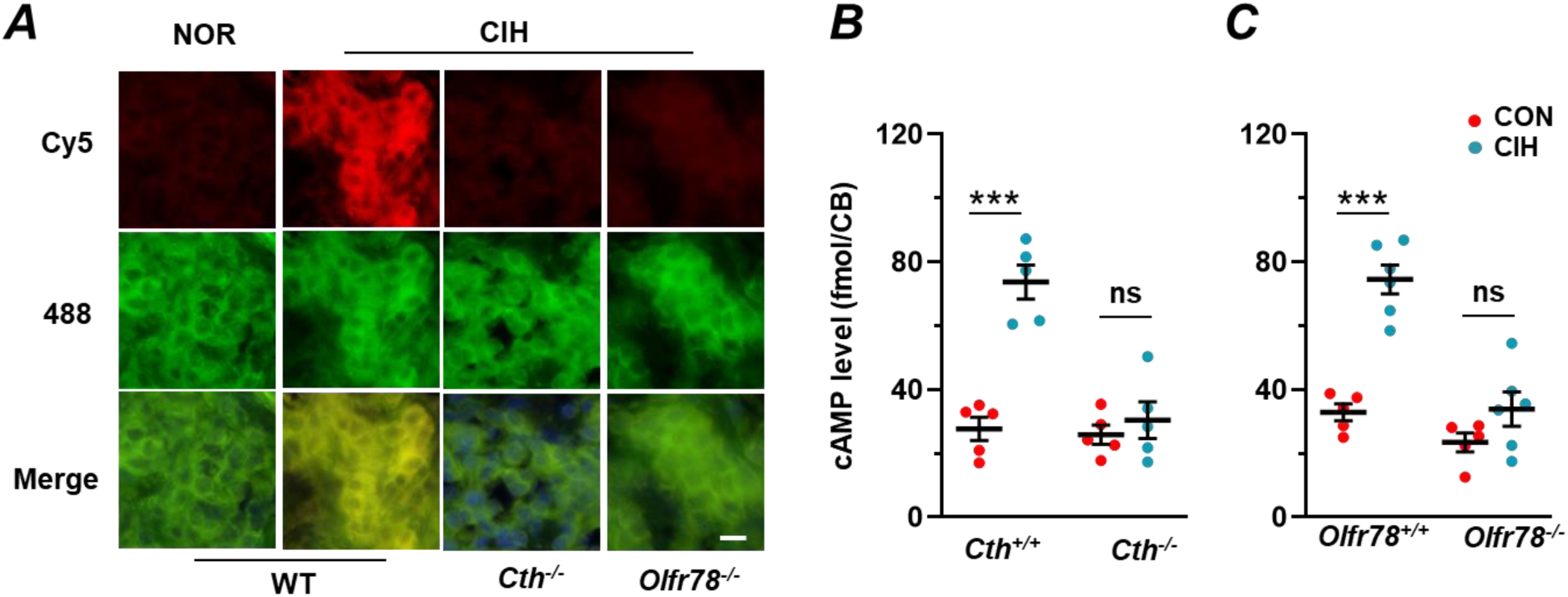
Chronic intermittent hypoxia (CIH) increases persulfidation in carotid bodies. **A**) Example of microscope images of persulfidation in carotid body sections from CIH and room air treated (NOR) wild type (WT), and *Cth* ^-/-^ and *Olfr78* ^-/-^ mice (n=3 mice with each genotype) treated with CIH. Cy5 signal represents persulfidation signal NBF-adducts recorded at 488 nm. CIH increased persulfidation in wildtype but not in *Cth* and *Olfr78* mutants. Scale bar, 10 µm. **B-C**. cAMP abundance was measured in carotid bodies from mice of the indicated genotypes after CIH treatment for 10d or room air (CON). Data are presented are individual data along with mean ± SEM (2 carotid bodies were pooled from one mouse for each experiment). *** *P* < 0. 001; ns, *P* > 0.05; two-way ANOVA followed by Holm-Sidak test.

### Carotid body (CB) sensory nerve response to CIH

We then examined the role of cAMP in CB responses to CIH. CIH has two effects on CB a) it enhances the CB sensory nerve (CSN) response to acute hypoxia and, b) induces a long-lasting increase in baseline CSN activity following acute intermittent hypoxia (AIH), known as sensory long-term facilitation (sensory LTF)(Peng *et al*., 2003). CSN activity was measured in CIH-treated wild-type (WT) and *Adcy3*-null mice. To eliminate the confounding influence of blood pressure changes in intact animals, CSN activity was recorded from an *ex vivo* CB preparation.

CIH treated wild-type mice showed enhanced CSN responses to hypoxia compared to room air controls (Fig. 3A). In contrast, CIH treated Adcy3 mutants showed absence of augmented CSN response to graded hypoxia (Fig. 3B). Furthermore, room air treated control *Adcy3* mutant mice showed reduced CSN response to hypoxia compared to wild-type controls (Fig. 3A and C). Morphometric analysis showed CB morphology in wild type was same as *Adcy3* mutants as indicated by glomus cell number and the ratio of glomus cells to CB volume (Fig. 3-figure supplement 1). Furthermore, CSN activation by sodium cyanide (NaCN 3µg/ml), a non-selective pharmacological activator of CB was similar in both *Adcy3* mutants and wild-type controls (Fig. 3-figure supplement 2).

**Figure 3.**
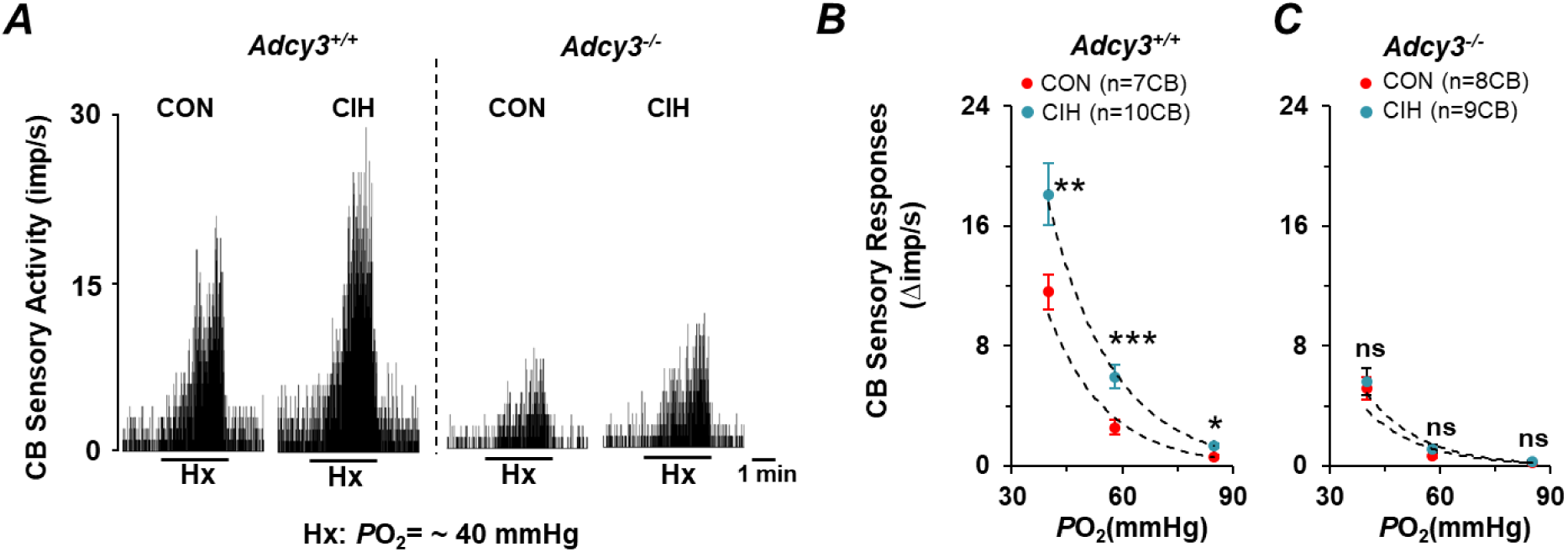
*Adcy3* null mice exhibit absence of CIH-induced augmented carotid body (CB) sensory response to hypoxia. **A**) Examples of CB sensory nerve response to hypoxia (medium P_O2_ ∼40 mmHg) in *Adcy3* ^+/+^ and *Adcy3* ^-/-^ mice treated with either room air (control = CON) or CIH. PO_2_ (mmHg) in the medium irrigating carotid bodies was measured by an O_2_ electrode placed close to the CB. Action potential frequencies are integrated and presented as impulses per second (imp/s). The duration of hypoxic challenge (Hx) is resented by black bars. B) Average data (mean ± SE) of CB sensory nerve responses to graded hypoxia presented as stimulus-evoked response minus baseline sensory nerve activity (delta impulse/s) in Adcy3 +/+ and Adcy3 -/- treated with either room air (CON) or CIH. Numbers in parenthesis represent the number of CBs in each group. ** and *** denote P < 0.01 and < 0.001, respectively and n.s. denotes P>0.05 analyzed by two-way ANOVA with repeated measures followed by Holm-Sidak test.

Sensory LTF was determined in CIH-treated wild-type and *Adcy3* mutant mice and room air treated controls. CIH treated wild-type mice showed a progressive increase in baseline CSN activity in response to five episodes of acute intermittent hypoxia (AIH; 30 seconds of hypoxia every 5 minutes), which persisted for an hour after terminating AIH (Fig. 4 A, C, and E), demonstrating the induction of sensory LTF by CIH. In contrast, CIH-induced sensory LTF was absent in *Adcy3* mutants (Fig. 4 B, D, and F).

**Figure 4.**
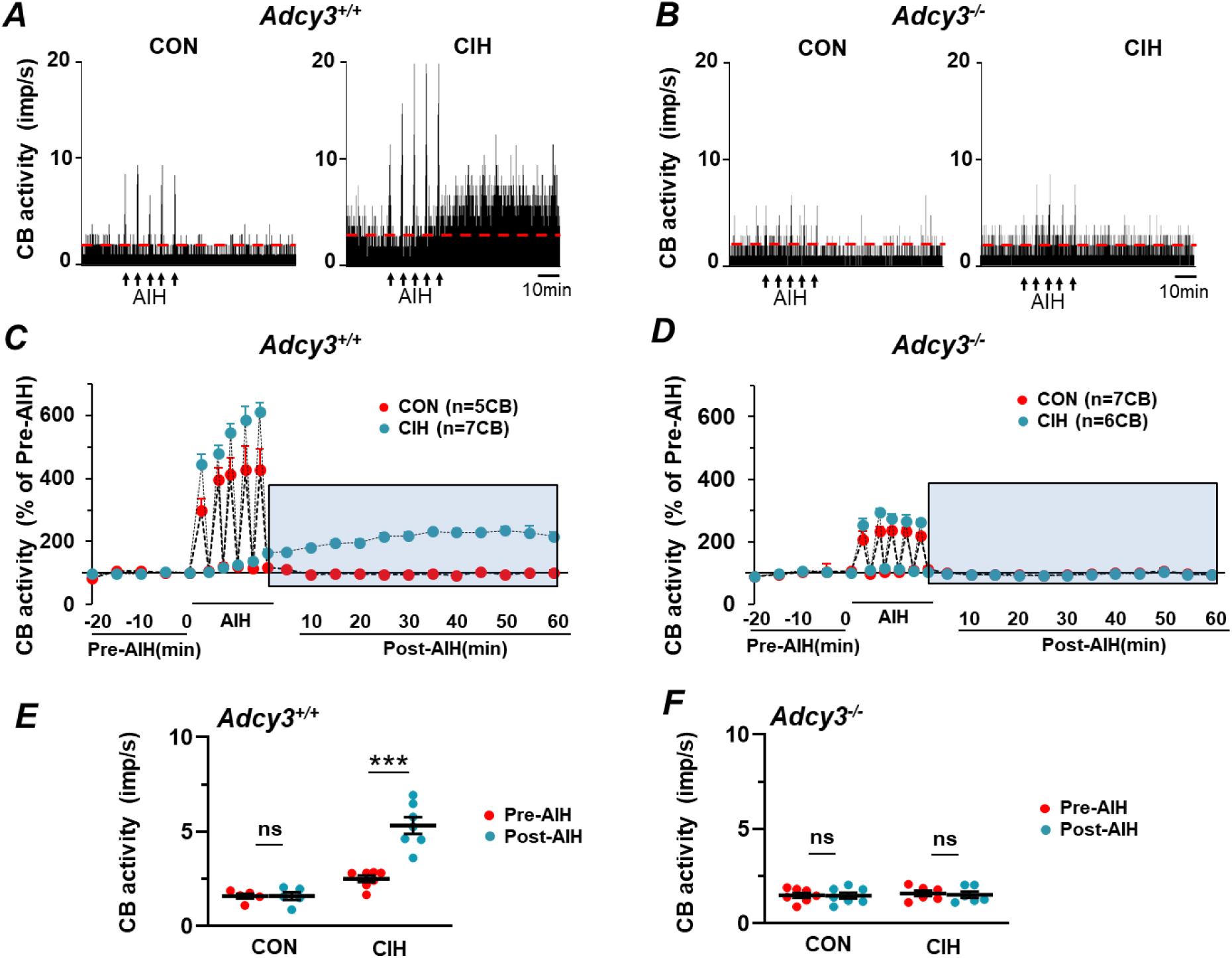
*Adcy3* null mice show absence of CIH-induced carotid body (CB) sensory long-term facilitation (sLTF). **A-B.** Examples of sLTF as measured by CB sensory nerve response to five episodes of acute intermittent hypoxia (AIH; denoted by arrows) in *Adcy3* ^+/+^ (A) and *Adcy3* ^-/-^ mice treated with either room air (CON) or CIH. Dashed red line represents the baseline activity. **C-D.** Average data (means ± SE) of CB sensory nerve activity before, during, and 60 min post-AIH presented as percentage of baseline activity (i.e., pre-AIH) in *Adcy3* ^+/+^ (C) and *Adcy3* ^-/-^ mice (D) treated with room air (CON) or CIH. Blue shaded are represent CB sensory activity during post-AIH. Numbers in parenthesis represent the number of CBs in each group. **E-F.** Individual data of CB sensory nerve activity before and during 60 min post-AIH along with means ± SE. in *Adcy3* ^+/-^ and *Adcy3* ^-/-^ mice treated with CIH. *** P<0.001; n.s. not significant P>0.05 compared to pre-AIH analyzed by two-way ANOVA with repeated measures followed by Holm-Sidak test.

### Cellular responses to CIH

Hypoxia increases Ca^2+^ influx in glomus cells, the primary O_2_ sensing cells of the CB (Kumar & Prabhakar, 2012). CIH augments Ca^2+^ influx by hypoxia (Makarenko *et al*., 2016). Glomus cell Ca^2+^ response to hypoxia was determined in CIH treated wild-type and *Adcy3* mutant mice. Cells from room air (controls) treated mice served as controls. CIH augmented Ca^2+^ influx by hypoxia in wild type but not in *Adcy3* mutant cells (Fig. 5 A-D). Control Ca ^2+^ response to hypoxia was reduced in *Adcy3* null glomus cells (Fig.5A). However, KCl (40mM)-evoked Ca^2+^ influx was comparable to wild type glomus cells (Fig. 5-figure supplement 1).

**Figure 5.**
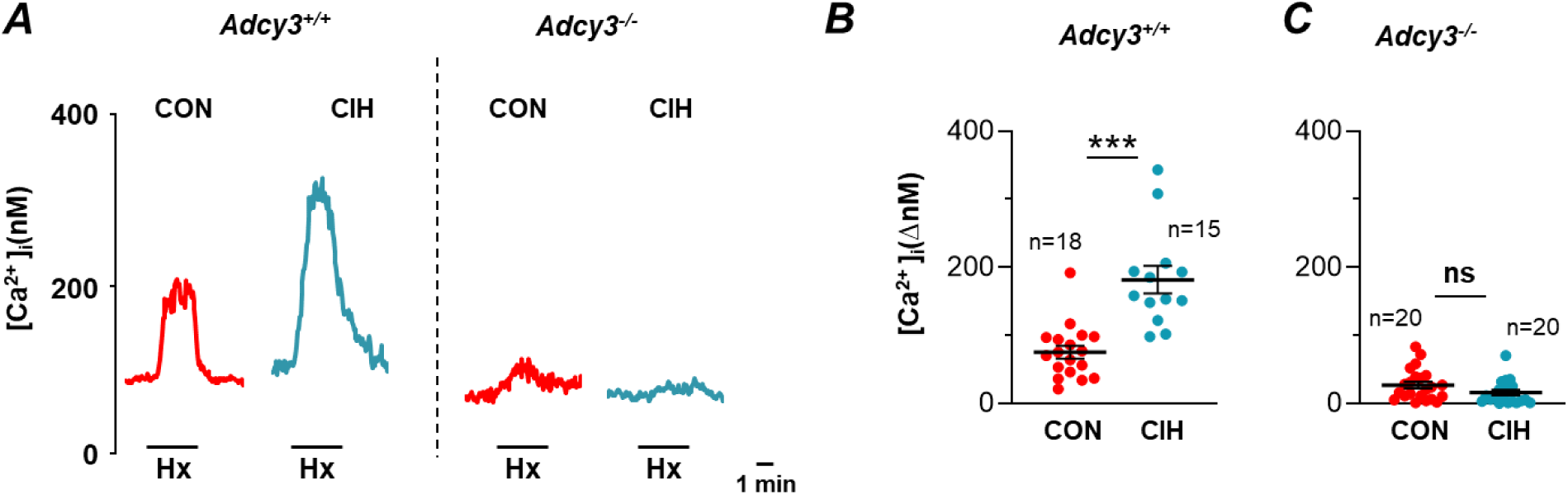
*Adcy3* mutant glomus cells exhibit impaired CIH-induced augmented Ca^2+^ response to hypoxia. **A.** Examples [Ca^2+^]_i_ responses of glomus cells to hypoxia (PO_2_ ∼40 mmHg, in *Adcy3^+/+^* and *Adcy3 ^-/-^* mice. **B-C.** Average (mean ± SEM) and individual data of [Ca^2+^]_i_ responses to hypoxia in *Adcy3 ^+/+^* (B) and *Adcy3 ^-/-^* (C) mice. ***, *P* < 0.001, n.s. not significant *P*>0.05 analyzed with Mann-Whitney Rank Sum Test.

### Cyclic nucleotide gated channel (CNG) mediates augmented Ca^2+^ influx by CIH

cAMP activates cyclic-nucleotide-gated (CNG) channels, which are nonselective cation channels. CNG channels are composed of α and β subunits. Glomus cells express the α2 subunit of CNG channels (Cnga2) (Peng *et al*., 2023). We tested the role of Cnga2 channels in augmented Ca^2+^ responses to CIH. Ca^2+^ responses to hypoxia were measured in glomus cells from CIH-treated wild-type and *Cnga2* heterozygous mice (*Cnga2*^+/-^ mice). Heterozygous *Cnga2* (*Cnga2*^+/-^) mice were studied due to the poor survival of *Cnga2* homozygous mice.

Glomus cells from CIH-treated wild-type mice showed augmented Ca^2+^ influx by hypoxia compared to controls (Fig. 6A-B). Although *Cnga2* mRNA was unaltered by CIH in wild type mice (Fig. 6-figure supplement 1), the augmented Ca^2+^ influx by CIH was absent in *Cnga2* mutant glomus cells (Fig. 6A and C). Ca^2+^ response to hypoxia was reduced in control*Cnga2* mutant cells (Fig. 6 A and C). However, KCl (40mM) evoked Ca^2+^ influx by was comparable in wild-type and *Cnga2* mutant cells (Fig. 6-figure supplement 2).

**Figure 6.**
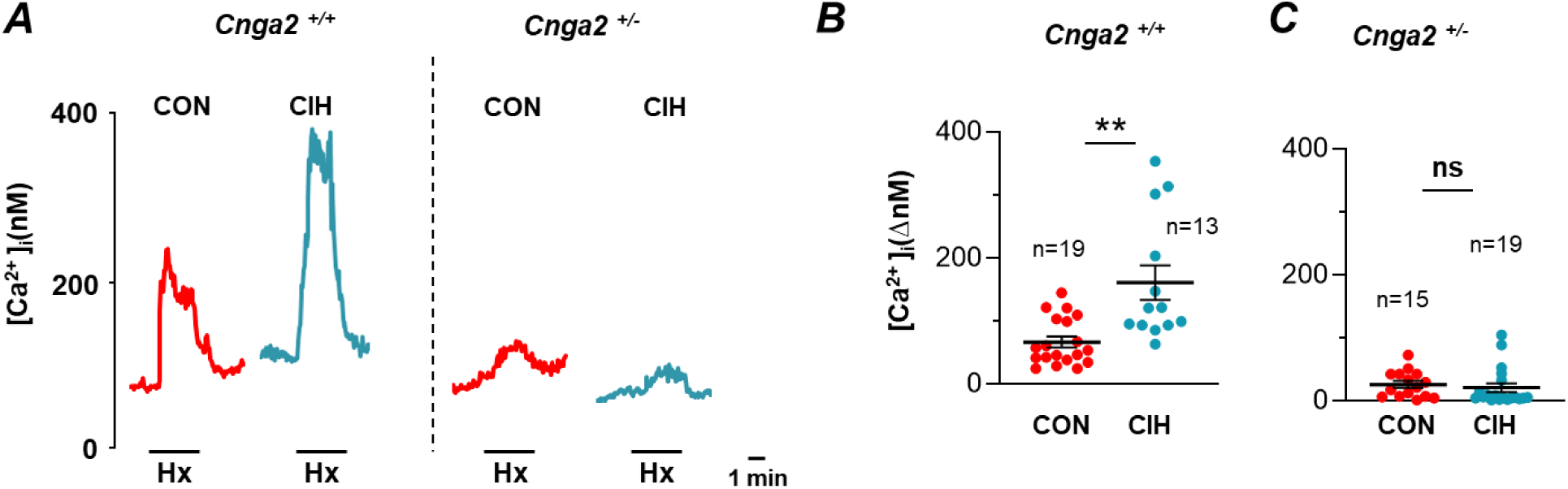
*Cnga2* mutant glomus cells exhibit impaired CIH-induced augmented Ca^2+^ response to hypoxia. **A.** Examples [Ca^2+^]_i_ responses of glomus cells to hypoxia (PO_2_ ∼40 mmHg, in *Cnga2^+/+^* and *Cnga2 ^+/-^* mice. **B-C.** Average (mean ± SEM) and individual data of [Ca^2+^]_i_ responses to hypoxia in *Cnga2^+/+^* (B) and *Cnga2 ^+/-^* (C) mice. **, *P* < 0.01, n.s. not significant *P*>0.05 analyzed with Mann-Whitney Rank Sum Test.

### Adcy3 null mice show impaired plasma catecholamine to CIH

CIH treated rodents show elevated splanchnic sympathetic nerve activity (SNA), and this response is absent in CB-ablated animals (Peng *et al*., 2014), suggesting that the CB chemoreflex is crucial for SNA activation by CIH. Given that *Adcy3* mutants showed absence of CB activation by CIH, we hypothesized that CIH-induced SNA should be absent in *Adcy3* mutants. Monitoring SNA in anesthetized mice was found to be technically challenging. Consequently, we measured plasma norepinephrine (NE) and epinephrine (Epi) levels as indices of SNA activation in wild- type and *Adcy3* mutant mice treated with CIH.

Examples of HPLC elution profiles of NE and Epi in plasma samples of normoxic and CIH-treated *Adcy3* ^+/+^ and *Adcy3* ^-/-^ mice are shown in Fig. 7A. NE and Epi were eluted at 6.9 and 8.1 minutes, respectively. Wild type (*Adcy3* ^+/+^) mice treated with CIH showed elevated plasma NE and Epi levels compared to room air-treated controls (Fig. 7A, B and C). In contrast, *Adcy3* mutant mice showed neither an increase in plasma NE nor Epi levels by CIH (Fig. 7A, B and C).

**Figure 7.**
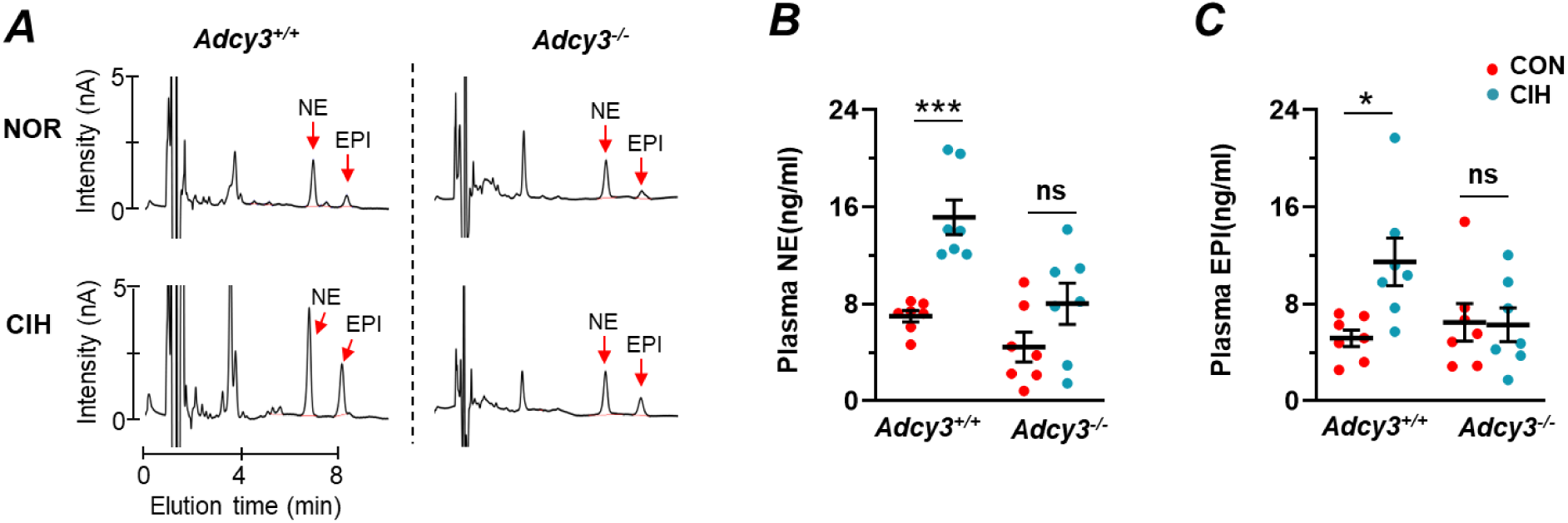
*Adcy3* null mice exhibit impaired elevation of circulating catecholamines by chronic intermittent hypoxia (CIH). **A.** Representative HPLC elution profiles of plasma norepinephrine (NE) and epinephrine (Epi) from *Adcy3 ^+/+^*and *Adcy3 ^-/-^* mice subjected to either room air (NOR) or 10d of CIH. The elution times for NE and EPI were 6.9 and 8.1 min, respectively. **B-C**. Average (mean ± SEM) and individual data of NE (**B**) and Epi (**C**) from both genotypes of mice subjected to room air (CON) and CIH. ***, *P* < 0.001; *, *P* < 0.05;n.s. not significant *P*>0.05 analyzed with two-way ANOVA followed by Holm-Sidak test.

### Adcy3 mutant mice exhibit absence of hypertension by CIH

CIH-treated rodents, like OSA patients, exhibit hypertension (Fletcher *et al*., 1992; Zoccal *et al*., 2007; Yuan *et al*., 2016; Peng *et al*., 2021). Blood pressures (BP) were measured in unsedated CIH-treated wild-type and *Adcy3* mutant mice using the tail-cuff approach. All measurements were taken between 11:00 AM and 1:00 PM to avoid the confounding influence of circadian variation. CIH-treated wild-type mice showed elevated systolic, diastolic, and mean blood pressure, and these responses were absent in *Adcy3*-mutant mice treated with CIH (Fig. 8).

**Figure 8.**
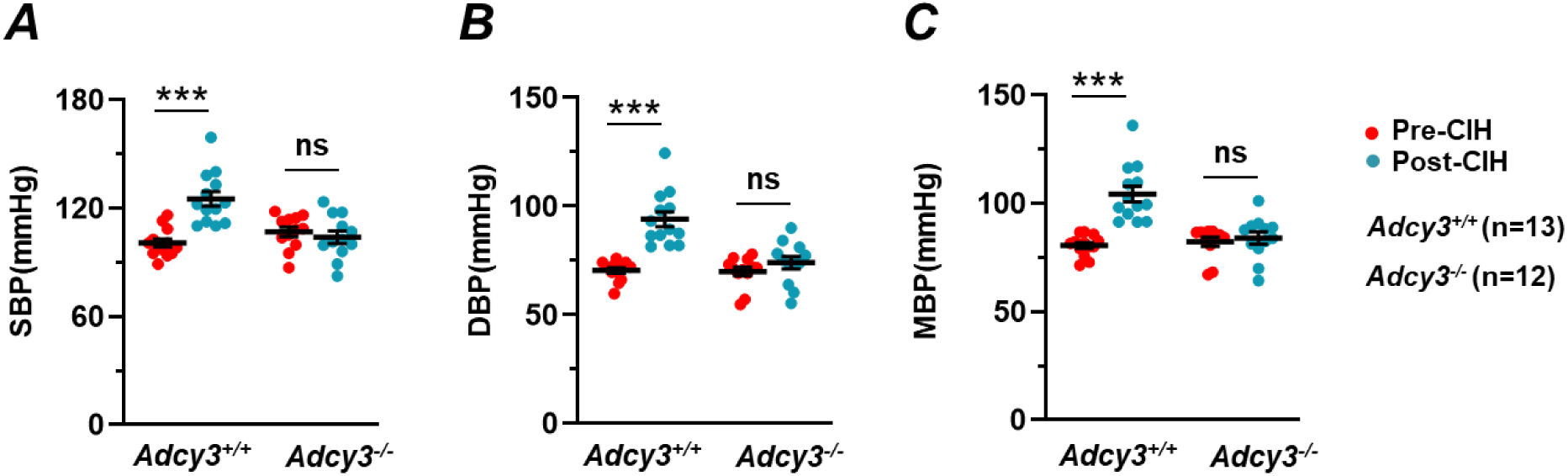
Chronic intermittent hypoxia-induced elevated blood pressure (BP) is absent in *Adcy3* null mice. Individual data points along with means ± SE of systolic (SBP) (A), diastolic (DBP) (B) and mean (MBP) (C) of Adcy3 +/- and Adcy3 -/- mice pre and post CIH. BP was monitored with tail-cuff method. Numbers represent the number of mice with each genotype. Shown are average (mean ± SEM) and individual data of BP of pre and post CIH of each genotype. ***, *P* < 0.001, n.s. not significant *P*>0.05 analyzed with two-way ANOVA with repeated measures followed by Holm-Sidak test.

## DISCUSSION

The present study establishes a previously uncharacterized role for cAMP in CB chemoreflex-mediated autonomic dysfunction induced by CIH, mimicking blood O_2_ profiles during obstructive sleep apnea (OSA). Our findings identify H_2_S-Olfr78 as the upstream signaling pathway for cAMP activation by CIH, and enhanced Ca^2+^ influx via cyclic nucleotide-gated channel (Cnga2) in glomus cells as the downstream signaling pathway to cAMP.

A major finding of this study is that CIH increases cAMP levels in the CB (Fig. 1). Adenylyl cyclases (Adcys) catalyze cAMP generation, and ten isoforms of Adcys have been identified in mammalian cells. Glomus cells of the CB express the Adcy3 isoform (Peng *et al*., 2023). Mice with targeted disruption of Adcy3 in glomus cells show a striking absence of increased cAMP in response to CIH, demonstrating that Adcy3 is the major isoform mediating cAMP elevation by CIH.

How might CIH increase cAMP? *Adcy3* mRNA levels in the carotid body remained unchanged by CIH, suggesting that increased cAMP levels are unlikely due to upregulation of the *Adcy3* gene. Adcy3 is associated with odorant receptor signaling (Su *et al*., 2009; Connelly *et al*., 2015). Murine CBs express a high abundance of the Olfr78 gene, which encodes the odorant receptor Olfr78 (Chang *et al*., 2015; Zhou *et al*., 2016). Hydrogen sulfide (H_2_S), derived from cystathionine-γ-lyase (CSE), activates Olfr78 through persulfidation of the Cys240 residue (Peng *et al*., 2023). CIH-treated CBs showed increased persulfidation, as indicated by an increased Cy5 signal, which was absent in either *Cth* or *Olfr78* mutant CBs (Fig. 2). These results suggest that CIH increases CSE-derived H_2_S, which in turn activates Olfr78 by persulfidation. Moreover, the CIH-induced increase in cAMP is absent in either *Cth* or *Olfr78* mutant CBs (Fig. 2). These results demonstrate that Adcy3 activation by CIH requires the H_2_S-Olfr78 pathway.

Chronic intermittent hypoxia (CIH) augmented the CB neural response to hypoxia and induced sensory long-term facilitation in wild-type mice, consistent with earlier reports (Yuan *et al*., 2016; Peng *et al*., 2021). In striking contrast, *Adcy3* mutant mice showed an absence of CB responses to CIH (Figs. 3 and 4). Control (room air treated) *Adcy3* mutant CBs exhibited reduced hypoxic sensitivity (Figs. 3 and 4). This impaired CB hypoxic sensitivity is unlikely due to altered CB morphology (Supplemental data, Fig. S2). Moreover, CSN stimulation by sodium cyanide (NaCN), a non-selective pharmacological activator of the CB, was similar in wild-type and *Adcy3* mutant mice (Fig. S3), indicating that the sensory nerve is functional in *Adcy3* mutants. The lack of CB activation by CIH is likely due to the absence of cAMP elevation in Adcy3 mutants. Consistent with this possibility, a cAMP analogue, like CIH, augmented the CB neural response to hypoxia similar to CIH (Peng *et al*., 2023).

Hypoxia-evoked Ca^2+^ influx in glomus cells is a crucial cellular event for CB neural activation (Kumar & Prabhakar, 2012). CIH augmented Ca^2+^ influx in wildtype but not in *Adcy3* mutant glomus cells, suggesting that cAMP mediates the augmented Ca^2+^ influx induced by CIH. How might cAMP increase Ca^2+^ influx? cAMP opens cyclic-nucleotide-gated channels, facilitating the influx of divalent cations such as Ca^2+^. Glomus cells express the Cnga2 protein (Peng *et al*., 2023). Although *Cnga2* mRNA levels remained unchanged by CIH, mice with partial deficiency of Cnga2 (*Cnga2*^+/-^) showed a striking absence of augmented Ca^2+^ influx induced by CIH, suggesting that cAMP is required for opening the Cnga2 channel. Besides cAMP, CNG channels can also be activated by cGMP. However, a cAMP analogue *augments*, whereas a cGMP analogue *inhibits*, the CB neural response to hypoxia (Peng *et al*., 2023). Further studies with direct monitoring of Cnga2 channel activity in glomus cells are needed to firmly establish the role of CNG channels in CIH. Notwithstanding this limitation, current results suggest that CNG, especially the Cnga2, mediates augmented Ca^2+^ influx, which is a downstream signaling event of cAMP activation by CIH.

Autonomic dysfunction, characterized by elevated sympathetic nerve activity and hypertension, is a major comorbidity in OSA patients (Somers *et al*., 1995; Nieto *et al*., 2000; Peppard *et al*., 2000; Dempsey *et al*., 2010). The autonomic dysfunction is mediated by CB chemoreflex (Hedner *et al*., 1992; Narkiewicz & Somers, 2003; Mansukhani *et al*., 2015). Consistent with earlier studies (Peng *et al*., 2006; Yuan *et al*., 2016; Peng *et al*., 2021), CIH-treated wild-type mice showed elevated sympathetic nerve activity, as evidenced by increased plasma norepinephrine (NE) and epinephrine (Epi) levels, as well as elevated systolic, diastolic, and mean blood pressures. In striking contrast, *Adcy3* mutant mice showed an absence of sympathetic nerve activation and hypertension, along with a lack of CB activation. These results suggest that CB activation involving cAMP signaling leads to sympathetic activation and hypertension in response to CIH.

Continuous positive airway pressure (CPAP) is the current treatment of choice for obstructive sleep apnea (OSA). However, not all patients comply with CPAP therapy, highlighting the need for alternative therapeutic strategies. Although CB ablation prevents autonomic dysfunction induced by chronic intermittent hypoxia (CIH) in experimental animals (Peng *et al*., 2014; Del Rio *et al*., 2016), it is not an ideal therapeutic intervention for alleviating the autonomic comorbidity of OSA, as the CB chemoreflex is vital for maintaining homeostasis under hypoxic conditions. Targeted ablation of *Adcy3* in glomus cells reduced, but did not abolish, the CB response to CIH. Therefore, reducing CB sensitivity to CIH with pharmacological blockade of cAMP generation by Adcy3 might offer a novel therapeutic strategy for treating autonomic dysfunction associated with OSA.

## MATERIAL AND METHODS

### General preparation

Experimental protocols were approved by the Institutional Animal Care and Use Committee of the University of Chicago (Protocol # ACUP 71811, approved on February 27, 2019). Studies were performed on age- and gender-matched adult wild-type (WT) mice (both males and females, 3-5 months old). The *TH*-Cre/*Adcy3*^f/f^ mice were obtained by crossing *Adcy3*^f/f^ mice (a gift from Drs. X. Chen, The University of New Hampshire, and D.R. Storm, The University of Washington) (Chen *et al*., 2016) with *TH*-Cre (JAX stock # 008601) at the University of Chicago. *Cth* null mice were obtained from Drs. S.H. Snyder (Johns Hopkins University School of Medicine and R. Wang, York University, Toronto, Ontario, Canada (Peng *et al*., 2010). *Cnga2*^+/-^ mice were sourced from JAX (stock # 002905). *Olfr78* null mice were initially generated by Bozza et al (Bozza *et al*., 2009) and subsequently backcrossed with C57BL/6 mice by Dr. Pluznick at The Johns Hopkins University School of Medicine, Baltimore, MD. All mice were on a C57BL/6 background. Experiments were performed by individuals blinded to the genotype.

### Exposure to Chronic Intermittent Hypoxia (CIH)

Mice housed in cages were placed in a specialized chamber for CIH treatment. The CIH paradigm consisted of 15 seconds of 5% inspired O2 (nadir) followed by 5 minutes of room air (normoxia), with 9 episodes per hour and 8 hours per day for 10 days, as previously described (Peng & Prabhakar, 2004; Peng *et al*., 2021). Oxygen and CO_2_ levels in the chamber were continuously monitored by an O_2_ analyzer (Alpha Omega Instrument; Series 9500), and CO_2_ levels were maintained between 0.2% and 0.5%. Mice treated with alternating cycles of room air instead of hypoxia in the same chamber served as controls.

### Measurement of carotid body sensory nerve activity

Carotid body sensory nerve (CSN) activity was recorded from *ex vivo* carotid bodies (CB) harvested 16 hours after terminating chronic intermittent hypoxia (CIH). The protocols for measuring CSN activity were the same as previously described (Peng *et al*., 2021; Peng *et al*., 2023). Briefly, carotid bodies along with the sinus nerves were harvested from urethane-anesthetized mice (urethane, 1.2 g/kg, i.p.). The tissues were placed in a 250 μL recording chamber and irrigated with warm physiological saline (35°C) at a rate of 3 mL/min. The composition of the physiological saline was (in mM): NaCl, 125; KCl, 5; CaCl2, 1.8; MgSO4, 2; NaH2PO4, 1.2; NaHCO3, 25; D-glucose, 10; sucrose, 5. The solution was bubbled with 21% O_2_/5% CO_2_. To facilitate the recording of action potentials, the sinus nerve was treated with 0.1% collagenase for 5 minutes. Action potentials (1–3 active units) were recorded from one of the nerve bundles using a suction electrode, filtered (bandpass, 100–3000 Hz), amplified (P511K; Grass Technologies, Natus Neurology, Middleton, WI), collected (sampling rate of 10 kHz), and stored on a computer with a data acquisition system (PowerLab/8P, AD Instruments, Colorado Springs, CO). ‘Single’ units were sorted based on the shape, height, and duration of the individual action potentials using the spike discrimination module. CBs were challenged with hypoxia by switching the perfusate equilibrated with desired amounts of O_2_, and the oxygen in the medium close to the CB was continuously monitored by a platinum electrode connected to a polarographic amplifier (Model 1900, A-M Systems, Sequim, WA).

### Sensory Long-Term Facilitation of the Carotid Body

Protocols for evoking sensory long-term facilitation (sLTF) of the carotid body (CB) were the same as previously described (Yuan *et al*., 2016; Peng *et al*., 2021). Briefly, baseline sensory activity was recorded for 20 minutes while irrigating the CB with room air-equilibrated medium (Medium PO2 ∼140 mmHg). This was followed by five episodes of intermittent hypoxia (Medium PO2 ∼40 mmHg; each hypoxic episode lasting 30 seconds) interspersed with 5 minutes of room air (normoxia). After completing the five episodes of intermittent hypoxia, sensory nerve activity was recorded continuously for 60 minutes while irrigating with room air-equilibrated medium.

### Primary glomus cell culture

The protocols for glomus cell isolation and maintenance of primary cultures are the same as previously described methods (Makarenko *et al*., 2012; Peng *et al*., 2020; Peng *et al*., 2023). Briefly, carotid bodies were harvested from mice anesthetized with urethane (1.2 g/kg, intraperitoneally). Glomus cells were dissociated using a mixture of collagenase P (2 mg/ml; Roche Applied Science, Indianapolis, IN), DNase (15 μg/ml; Sigma-Aldrich), and BSA (3 mg/ml; Sigma-Aldrich) at 37°C for 20 minutes. This was followed by a 15-minute incubation in Locke’s buffer containing DNase (30 μg/ml). The cells were then plated on collagen (type VII; Sigma-Aldrich)-coated coverslips and maintained at 37°C in an incubator with 7% CO2 and 20% O2 for 12–18 hours. The growth medium consisted of DMEM/F-12 medium (Invitrogen, Thermo Fisher Scientific, Waltham, MA), supplemented with 1% fetal bovine serum (FBS), insulin-transferrin-selenium (ITS-X; Invitrogen), and a 1% penicillin-streptomycin-glutamine mixture (Invitrogen).

### Measurements of [Ca^2+^]_i_

A coverslip with glomus cells was incubated in Hanks’ balanced salt solution (HBSS, Thermo Fisher Scientific) containing 2 μM fura-2 AM (Biotium, Inc., Fremont, CA) and 1 mg/mL BSA for 30 minutes. It was then washed in a fura-2-free solution for another 30 minutes at 37°C. The coverslip was transferred to a chamber for measuring intracellular calcium concentration ([Ca^2+^]i). Background fluorescence was obtained at 340- and 380-nm wavelengths from an area of the coverslip devoid of cells. On each coverslip, glomus cells were identified by their characteristic clustering, and individual cells were imaged using a Leica microscope equipped with a Hamamatsu camera (model C11440) and the HC Image software (version 4.5.1.3). Image pairs (one at 340 nm and the other at 380 nm) were obtained every 2 seconds by averaging 16 frames at each wavelength. Data were continuously collected throughout the experiment. Background fluorescence was subtracted from the cell data obtained at each wavelength.

Fluorescence intensity was calculated by dividing the image obtained at 340 nm by the image collected at 380 nm to obtain a ratiometric image. Ratios were converted to free [Ca^2+^]i using calibration curves constructed in vitro by adding fura-2 (50 μM free acid) to solutions containing known concentrations of Ca^2+^ (0 – 2,000 nM). The recording chamber was continually irrigated with warm physiological saline (31°C) from gravity-fed reservoirs. The composition of the saline was the same as that used for recording CB sensory nerve activity.

### Measurements of cAMP in carotid bodies

Carotid bodies were harvested from anesthetized mice (urethane, 1.2 g/kg, i.p.) treated with either CIH or room air. The cAMP abundance in the carotid bodies was measured using a cAMP ELISA kit (STA-501, Cell Biolabs, Inc., San Diego, CA) according to the manufacturer’s instructions. In each experiment, two carotid bodies were pooled, homogenized in 100 μL lysis buffer on ice for 30 minutes, and centrifuged for 5 minutes at 16,000 × g. The supernatant (50 μL) was added to a well of a 96-well plate. Diluted peroxidase cAMP tracer conjugate (25 μL) and diluted rabbit anti-cAMP polyclonal antibody (50 μL) were added to each well, and the plate was incubated for 30 minutes at room temperature. After washing five times with wash buffer, each well was incubated with 100 μL Chemiluminescent reagent for 5 minutes, and luminescence was read with a microplate luminometer. For each experiment, a corresponding standard curve was generated with cAMP standards, and this standard curve was used to calculate the cAMP content in the carotid bodies. The cAMP content is expressed as femtomoles of cAMP per carotid body. The detection limit of the assay was 1 pM of cAMP.

### Carotid body morphology

Mice anesthetized with urethane were intra-cardially perfused with heparinized PBS (pH 7.4) at a rate of 10 ml/min for 10 minutes, followed by buffered formaldehyde (4% formalin; Fischer Scientific) for 30 minutes. Carotid bifurcations were removed and placed in 4°C 4% paraformaldehyde–PBS for 1 hour. After washing with PBS, the carotid bifurcations were placed in 30% sucrose–PBS at 4°C for 24 hours. Specimens were frozen in Tissue Tek (OCT; VWR Scientific), serially sectioned at 8 μm (Leica CM1900), and mounted on collagen-coated coverslips. Sections were blocked in PBS containing 1% normal goat serum and 0.2% Triton X- 100, and then incubated with monoclonal rabbit anti-tyrosine hydroxylase (TH) antibody (Pel-Freez Biologicals; #P4010-0; dilution 1:4,000), followed by five washes with PBS containing 0.05% Triton X-100. Antibody binding was detected using Texas red-conjugated goat anti-mouse IgG diluted 1:250 in PBS containing 1% normal goat serum and 0.2% Triton X-100 (1 hour at 37°C) and washed with PBS containing 0.05% Triton X-100 (five times). Sections were mounted in Vectashield with DAPI (Vector Laboratories; #H-1200) and visualized using a using an all-in-one fluorescent microscope (BZ-X810; Keyence Corp. of America, Itasca, IL).

### Monitoring persulfidation in the carotid body by microscopy

Protocols for detecting persulfidation in carotid body sections are the same as described earlier (Zivanovic *et al*., 2019; Peng *et al*., 2023) . Carotid bifurcations were removed from anesthetized mice treated either with CIH or room air (controls). Tissues were fixed in 100% methanol at -20°C for 4 hours. The methanol-fixed tissue was placed in 30% sucrose–PBS overnight at 4°C, then frozen in Tissue Tek (OCT; VWR Scientific), serially sectioned at 8 μm (Leica CM1900), and mounted on collagen-coated coverslips. Tissue sections were treated with acetone for 5 minutes at -20°C, followed by three washes with PBS for 5 minutes at 37°C. Following incubation with 1 mM NBF-Cl (4-Chloro-7-nitrobenzofurazan, Sigma-Aldrich, #163260) in PBS for 2 hours at 37°C, sections were washed with PBS at room temperature, left overnight, and then washed at 4°C with agitation. Sections were then incubated with 10 µM DAz-2: Cy-5 mix (DAz-2, Cayman Chemical, #13382; Cy-5, Lumiprobe, #B30B0) in PBS for 30 minutes at 37°C. As negative controls, sections were incubated with 10 mM DAz-2: Cy-5 click mix prepared without DAz-2. Subsequently, sections were washed with PBS overnight with agitation, protected from light, followed by three washes with 100% methanol for 10 minutes at room temperature, and five washes with PBS for 5 minutes. Sections were mounted in Vectashield with DAPI (Vector Laboratories *H-1200) and visualized using an all-in-one fluorescent microscope (BZ-X810; Keyence Corp. of America, Itasca, IL). Sections were examined at 488 nm (for NBF-adducts) and 633 nm (Cy5 for PSSH).

### Genotyping of *Th*-Cre+/*Adcy3 ^fl/fl^* (referred as *Adcy3*^-/-^)mice

A tail biopsy (∼2 mm long) was collected from mice anesthetized with isoflurane. Genomic DNA was extracted, and polymerase chain reaction (PCR) was performed to amplify the following primer sets targeting WT, TH-Cre, and Adcy3 floxed alleles. Genotyping of *Adcy3* flox mice was conducted using protocols from Dr. X. Chen (The University of New Hampshire). The primers used were: *Adcy3* flox forward: ACC CTT TGA GGC CAG GGG CAA; *Adcy3*3 flox reverse: CTG CGG TGA GAG CCT GGC ACA; Both primers were used in a single reaction, with expected bands for wild type and Adcy3 flox at 102 bp and 200 bp, respectively.

Genotyping of *TH*-Cre mice was performed using protocol #21729 provided by the Jackson Laboratory. The primers used were: Transgene forward: GAG ACA GAA CTC GGG ACC AC Transgene reverse: AGG CAA ATT TTG GTG TAC GG; Internal Positive Control Forward: AGT GGC CTC TTC CAG AAA TG; Internal Positive Control Reverse: TGC GAC TGT GTC TGA TTT CC. All four primers were used in a single reaction, with expected bands for the transgene and internal positive control at 300 bp and 521 bp, respectively. *Adcy3*^-/-^ mice exhibited both *TH*- Cre and *Adcy3* flox/flox expression.

### Measurements of mRNA by Quantitative Real-Time PCR

Real-time polymerase chain reaction (RT-PCR) assay was performed using a MiniOpticon system (Bio-Rad Laboratories) with SYBR Green ER two-step qRT-PCR kit (No. 11764-100, Invitrogen) as described (Peng *et al*., 2009). Briefly, carotid bodies were harvested from anesthetized mice, and RNA was extracted using TRIZOL reagent and reverse transcribed using iScript reverse transcriptase super mix (Bio-Rad). The mRNA abundance was calculated with the comparative threshold (CT) method using the formula “2−^ΔCT^” where ΔCT is the difference between the threshold cycle of the given target cDNA expressed in normoia and CIH carotid bodies. The CT value was taken as a fractional cycle number at which the emitted fluorescence of the sample passes a fixed threshold above the baseline. Values were compared with the internal standard the *18S* gene. Purity and specificity of all products were confirmed by omitting the template and performing a standard melting curve analysis. The following primers were used: *18s:* Forward Primer: CGC CGC TAG AGG TGA AAT TC; Reverse Primer: CGA ACC TCC GAC TTT CGT TCT; *Adcy3:* Forward Primer: TCA TCG TGG GCA TCA TGT CCT A; Reverse primer: TGC TCC TCC AGA TTC ATC TTC ACC; *Cnga2*: Forward primer:GCC TGC TTC AGT GAT CTA CAG AG; Reverse Primer: TTC TAG GAA GCC TGT GCG CA.

### Blood Pressure Measurements

Blood pressure (BP) was measured using the tail-cuff method in unanesthetized mice with a noninvasive BP system (IITC Life Science Inc, Woodland Hills, CA) as previously described (Peng *et al*., 2021). To minimize variations in BP measurements, the following measures were taken: a) Mice were allowed to acclimate to the recording apparatus for at least 1 hour for 3 consecutive days. b) BP was measured in the same mouse before and after CIH, with each mouse serving as its own control. A minimum of five measurements were taken for each mouse. All measurements were conducted between 9:00 AM and 11:00 AM, 16 hours after terminating CIH, to exclude confounding influences from circadian variations.

### Measurements of plasma catecholamines

Blood samples (∼300 μL) were collected from anesthetized mice (urethane, 1.2 g/kg ip) by cardiac puncture and placed in heparinized (30 U/mL of blood) ice-cold microcentrifuge tubes. Plasma was separated by centrifugation and stored at -80°C. Plasma norepinephrine (NE) levels were determined by high-pressure liquid chromatography combined with electrochemical detection (HPLC-ECD) using dihydroxybenzylamine (DBA) as an internal standard, as previously described (Kumar *et al*., 2006). NE and Epi levels were corrected for recovery loss and expressed as nanograms of NE or Epi per 1 mL of plasma.

### Statistical analysis of the data

Carotid body sensory activity (CSN activity from ‘single’ units) was averaged for 3 min prior to hypoxic challenge and during the entire 3 min of hypoxic challenge and expressed as impulses per second unless otherwise stated. BP measurements include systolic, diastolic, and mean BP. All data are presented as individual data points along with mean ± SEM, unless otherwise stated. The following statistical methods were employed. A t-test was performed for the data with normal distribution (Shapiro-WilK test) and equal variances (Levene’s Median test). Otherwise, Mann- Whitney Rank Sum test was performed. To determine whether the means of two or more groups are affected by two different factors (genotype/treatment/time point), two-way ANOVA, or two- way ANOVA with repeated measures was performed. All statistical analysis was performed using Sigma Plot (version 11) and P values <0.05 were considered significant.

## Acknowledgement

The authors thank Drs. S. H. Snyder and R. Wang for providing *Cth* null mice; Dr. Pluznick at The Johns Hopkins for *Olfr78* null mice; Drs. X. Chen, The University of New Hampshire, and D.R. Storm, The University of Washington for *Adcy3 ^fl/fl^* mice.

## Author contributions

N.R.P. conceived the project and wrote the manuscript; Y-J.P. performed neurophysiology, and measured [Ca^2+^]_i;_ M.H. and Y-J.P. performed blood pressure measurements. N.W., and J.N., performed persulfidation, histochemistry and mRNA measurements; X.S. and M.H. measured plasma catecholamines with HPLC; MH performed genotyping.

## Data and materials availability

All data, and materials used in the analysis are available in some form to any researcher for purposes of reproducing or extending the analysis.

---

**Figure 1-figure supplement 1.**
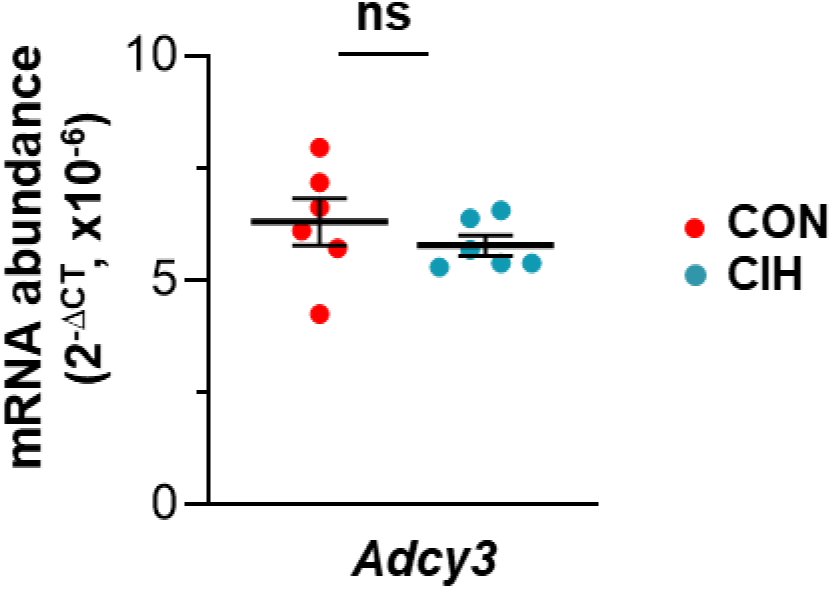
*Adcy3* mRNA in the carotid body was unaltered by chronic intermittent hypoxia (CIH). *Adcy3* mRNA was analyzed by RT-PCR assay in carotid bodies of BL6 mice subjected to either room air (CON) or 10d of CIH. n.s. denotes P>0.05 analyzed with t- test.

**Figure 3-figure supplement 1.**
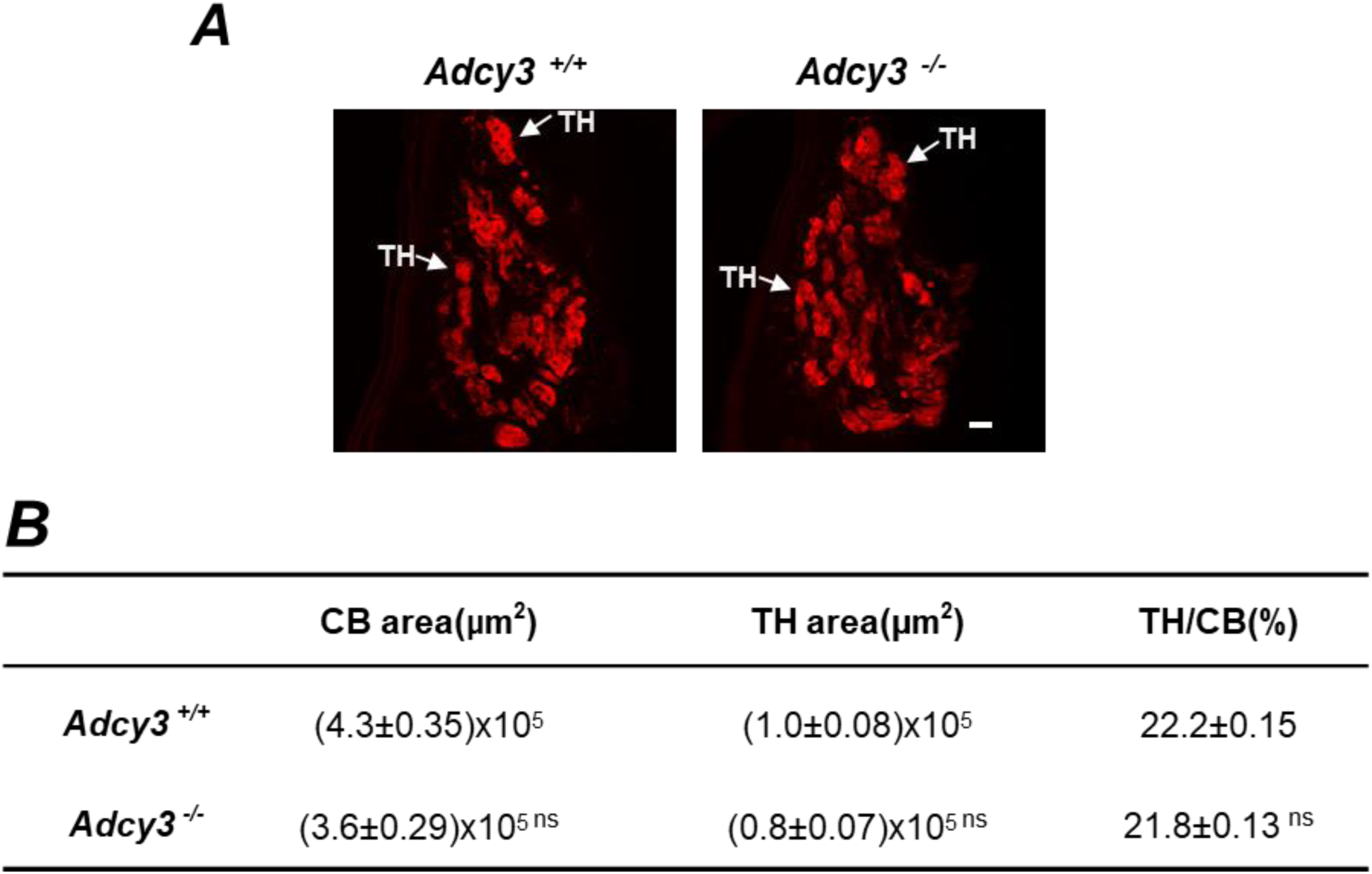
Morphometric analysis of carotid bodies of *Adcy3* ^+/+^ and *Adcy3* ^-/-^ mice. **(A)** Carotid body sections were stained with antibodies specific for tyrosine hydroxylase (TH). Scale bar, 20 µm. (B) Data were analyzed for CB area, TH positive cell area and ratio of TH/CB area. All data were presented as mean ±SEM from 4 mice of each genotype. n.s. not significant P>0.05 *Adcy3* ^-/-^ compared to *Adcy3* ^+/+^ mice, analyzed with t-test.

**Figure 3-figure supplement 2.**
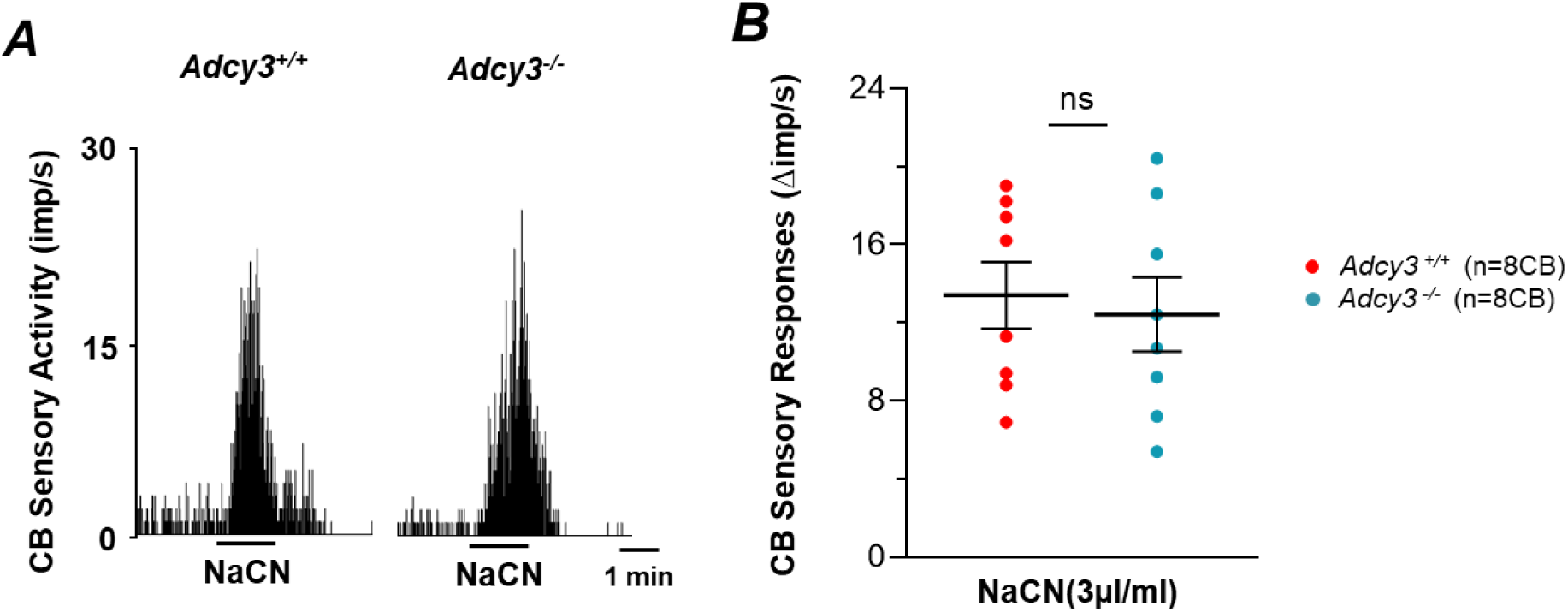
Carotid body (CB) sensory nerve (CSN) responses to sodium cyanide (NaCN) are comparable between *Adcy3* ^+/+^ and *Adcy3* ^-/-^ mice. **A.** Example of CSN response to NaCN (3µg/ml) in *Adcy3* ^+/+^ and *Adcy3* ^-/-^mice. **B**. Average (mean ±SEM) and individual data points of CSN response (NaCN-baseline delta imp/sec) are shown. Numbers in parentheses indicate number of CBs. n.s. not significant P>0.05 analyzed with t-test.

**Figure 5-figure supplement 1.**
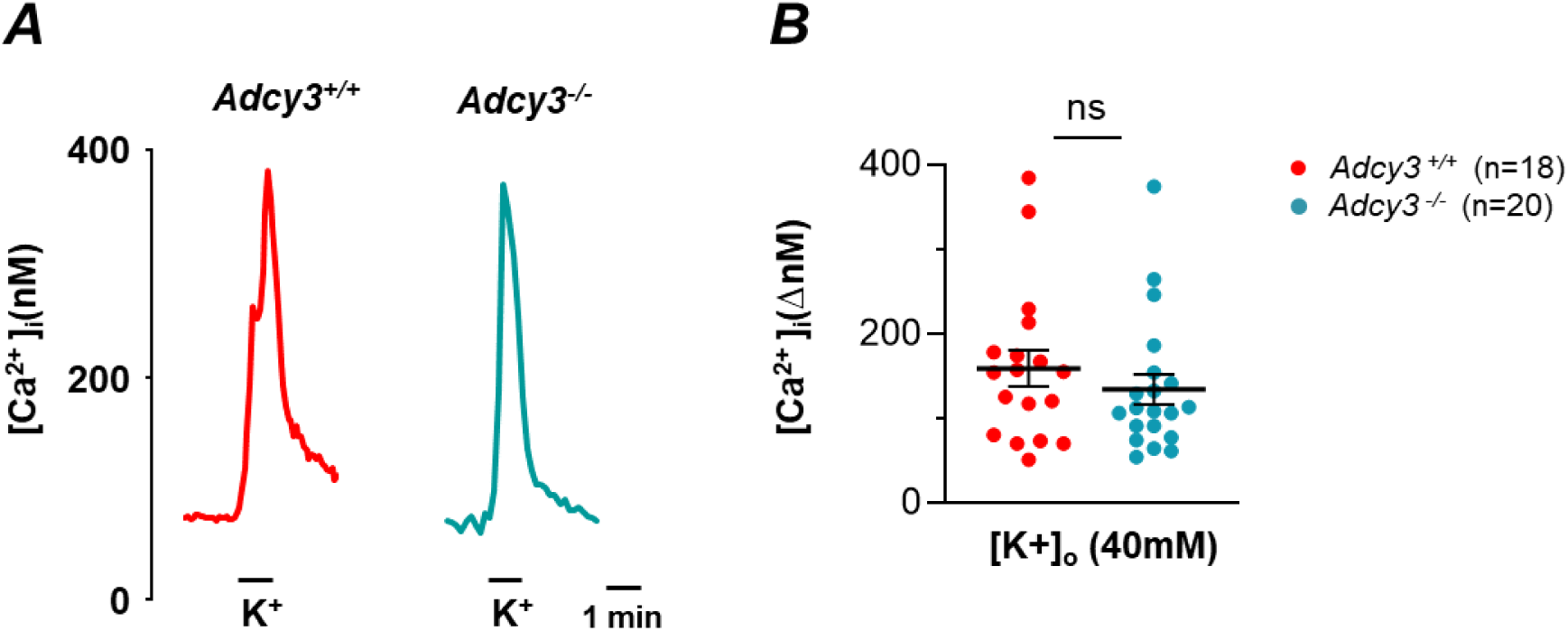
Comparable [Ca^2+^]_i_ responses of *Adcy3 ^+/+^* and Adcy3 ^-/-^ glomus cells to potassium chloride (KCl 40mM). **A**. Examples of [Ca^2+^]_i_ responses of glomus cells in genotypes of both mice. **B.** Average (mean ±SEM) and individual data points of Ca^2+^ responses presented as KCl-baseline (delta imp/sec). Numbers in parentheses indicate number of cells. n.s. not significant P>0.05 analyzed with Mann-Whitney Rank Sum Test.

**Figure 6-figure supplement 1.**
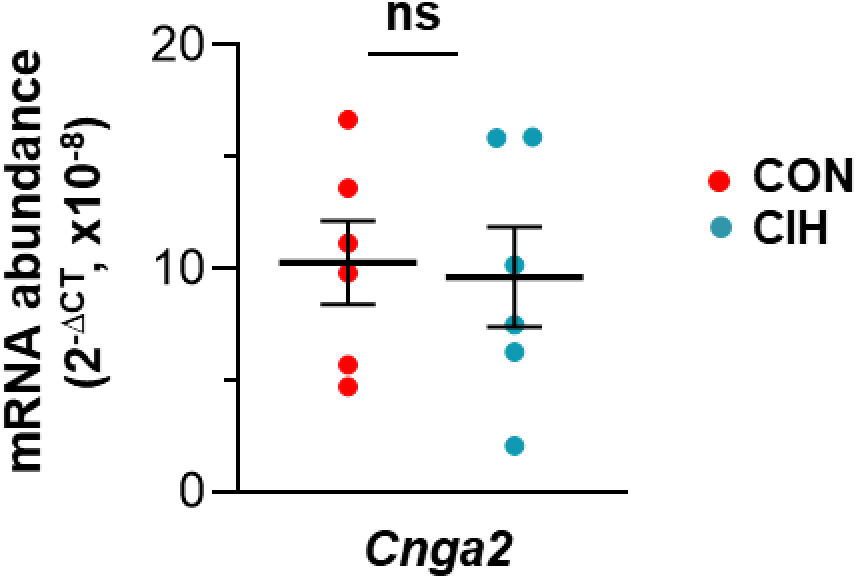
*Cnga2* mRNA in the carotid body was unaltered by chronic intermittent hypoxia (CIH). *Cnga2* mRNA was analyzed by RT-PCR assay in carotid bodies from BL6 wild type subjected to room air (CON) or 10d of CIH. n.s. denotes P>0.05 analyzed with t-test.

**Figure 6-figure supplement 2.**
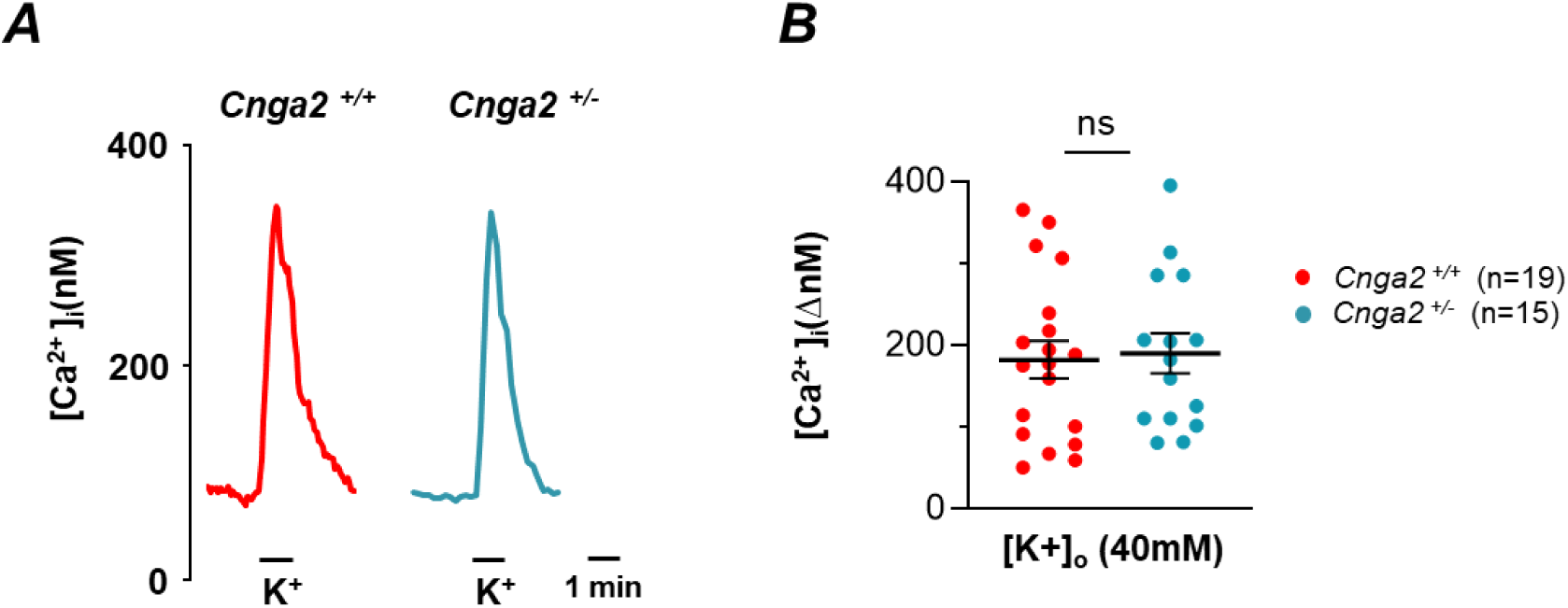
Comparable [Ca^2+^]_i_ responses of *Cnga2 ^+/+^* and Cnga2 ^+/-^ glomus cells to potassium chloride (KCl 40mM). **A**. Examples of [Ca^2+^]_I_ responses of glomus cells of genotypes. **B.** Average (mean ±SEM) and individual data points of Ca^2+^ responses presented as KCl-baseline (delta imp/sec). Numbers in parentheses indicate number of cells. n.s. not significant P>0.05 analyzed with Mann-Whitney Rank Sum Test.

## REFERENCES

1. Bozza T, Vassalli A, Fuss S, Zhang JJ, Weiland B, Pacifico R, Feinstein P & Mombaerts P. (2009). Mapping of class I and class II odorant receptors to glomerular domains by two distinct types of olfactory sensory neurons in the mouse. Neuron 61, 220–233.

2. Chang AJ, Ortega FE, Riegler J, Madison DV & Krasnow MA. (2015). Oxygen regulation of breathing through an olfactory receptor activated by lactate. Nature 527, 240–244.

3. Chen X, Luo J, Leng Y, Yang Y, Zweifel LS, Palmiter RD & Storm DR. (2016). Ablation of Type III Adenylyl Cyclase in Mice Causes Reduced Neuronal Activity, Altered Sleep Pattern, and Depression-like Phenotypes. Biol Psychiatry 80, 836–848.

4. Connelly T, Yu Y, Grosmaitre X, Wang J, Santarelli LC, Savigner A, Qiao X, Wang Z, Storm DR & Ma M. (2015). G protein-coupled odorant receptors underlie mechanosensitivity in mammalian olfactory sensory neurons. Proc Natl Acad Sci U S A 112, 590–595.

5. Del Rio R, Andrade DC, Lucero C, Arias P & Iturriaga R. (2016). Carotid Body Ablation Abrogates Hypertension and Autonomic Alterations Induced by Intermittent Hypoxia in Rats. Hypertension 68, 436–445.

6. Dempsey JA, Veasey SC, Morgan BJ & O’Donnell CP. (2010). Pathophysiology of sleep apnea. Physiol Rev 90, 47–112.

7. Fletcher EC, Lesske J, Behm R, Miller CC, 3rd, Stauss H & Unger T. (1992). Carotid chemoreceptors, systemic blood pressure, and chronic episodic hypoxia mimicking sleep apnea. J Appl Physiol 72, 1978–1984.

8. Hedner JA, Wilcox I, Laks L, Grunstein RR & Sullivan CE. (1992). A specific and potent pressor effect of hypoxia in patients with sleep apnea. Am Rev Respir Dis 146, 1240–1245.

9. Kumar GK, Rai V, Sharma SD, Ramakrishnan DP, Peng YJ, Souvannakitti D & Prabhakar NR. (2006). Chronic intermittent hypoxia induces hypoxia-evoked catecholamine efflux in adult rat adrenal medulla via oxidative stress. J Physiol 575, 229–239.

10. Kumar P & Prabhakar NR. (2012). Peripheral chemoreceptors: function and plasticity of the carotid body. Compr Physiol 2, 141–219.

11. Makarenko VV, Ahmmed GU, Peng YJ, Khan SA, Nanduri J, Kumar GK, Fox AP & Prabhakar NR. (2016). CaV3.2 T-type Ca2+ channels mediate the augmented calcium influx in carotid body glomus cells by chronic intermittent hypoxia. J Neurophysiol 115, 345–354.

12. Makarenko VV, Nanduri J, Raghuraman G, Fox AP, Gadalla MM, Kumar GK, Snyder SH & Prabhakar NR. (2012). Endogenous H2S is required for hypoxic sensing by carotid body glomus cells. Am J Physiol Cell Physiol 303, C916–923.

13. Mansukhani MP, Wang S & Somers VK. (2015). Chemoreflex physiology and implications for sleep apnoea: insights from studies in humans. Exp Physiol 100, 130–135.

14. Narkiewicz K & Somers VK. (2003). Sympathetic nerve activity in obstructive sleep apnoea. Acta Physiol Scand 177, 385–390.

15. Nieto FJ, Young TB, Lind BK, Shahar E, Samet JM, Redline S, D’Agostino RB, Newman AB, Lebowitz MD & Pickering TG. (2000). Association of sleep-disordered breathing, sleep apnea, and hypertension in a large community-based study. Sleep Heart Health Study. JAMA 283, 1829–1836.

16. Peng Y-J, Nanduri J, Raghuraman G, Souvannakitti D, Gadalla MM, Kumar GK, Snyder SH & Prabhakar NR. (2010). H2S mediates O-2 sensing in the carotid body. Proc Natl Acad Sci U S A 107, 10719–10724.

17. Peng YJ, Gridina A, Wang B, Nanduri J, Fox AP & Prabhakar NR. (2020). Olfactory receptor 78 participates in carotid body response to a wide range of low O_2_ levels but not severe hypoxia. J Neurophysiol 123, 1886–1895.

18. Peng YJ, Nanduri J, Wang N, Kumar GK, Bindokas V, Paul BD, Chen X, Fox AP, Vignane T, Filipovic MR & Prabhakar NR. (2023). Hypoxia sensing requires H2S-dependent persulfidation of olfactory receptor 78. Sci Adv 9, eadf3026.

19. Peng YJ, Nanduri J, Yuan G, Wang N, Deneris E, Pendyala S, Natarajan V, Kumar GK & Prabhakar NR. (2009). NADPH oxidase is required for the sensory plasticity of the carotid body by chronic intermittent hypoxia. J Neurosci 29, 4903–4910.

20. Peng YJ, Overholt JL, Kline D, Kumar GK & Prabhakar NR. (2003). Induction of sensory long- term facilitation in the carotid body by intermittent hypoxia: implications for recurrent apneas. Proc Natl Acad Sci U S A 100, 10073–10078.

21. Peng YJ & Prabhakar NR. (2004). Effect of two paradigms of chronic intermittent hypoxia on carotid body sensory activity. J Appl Physiol 96, 1236–1242; discussion 1196.

22. Peng YJ, Su X, Wang B, Matthews T, Nanduri J & Prabhakar NR. (2021). Role of olfactory receptor78 in carotid body-dependent sympathetic activation and hypertension in murine models of chronic intermittent hypoxia. J Neurophysiol 125, 2054–2067.

23. Peng YJ, Yuan G, Khan S, Nanduri J, Makarenko VV, Reddy VD, Vasavda C, Kumar GK, Semenza GL & Prabhakar NR. (2014). Regulation of hypoxia-inducible factor-α isoforms and redox state by carotid body neural activity in rats. J Physiol 592, 3841–3858.

24. Peng YJ, Yuan G, Ramakrishnan D, Sharma SD, Bosch-Marce M, Kumar GK, Semenza GL & Prabhakar NR. (2006). Heterozygous HIF-1alpha deficiency impairs carotid body- mediated systemic responses and reactive oxygen species generation in mice exposed to intermittent hypoxia. J Physiol 577, 705–716.

25. Peppard PE, Young T, Palta M & Skatrud J. (2000). Prospective study of the association between sleep-disordered breathing and hypertension. N Engl J Med 342, 1378–1384.

26. Prabhakar NR. (2013). Sensing hypoxia: physiology, genetics and epigenetics. J Physiol 591, 2245–2257.

27. Redline S, Azarbarzin A & Peker Y. (2023). Obstructive sleep apnoea heterogeneity and cardiovascular disease. Nat Rev Cardiol 20, 560–573.

28. Somers VK & Abboud FM. (1993). Chemoreflexes--responses, interactions and implications for sleep apnea. Sleep 16, S30–33; discussion S33-34.

29. Somers VK, Dyken ME, Clary MP & Abboud FM. (1995). Sympathetic neural mechanisms in obstructive sleep apnea. J Clin Invest 96, 1897–1904.

30. Su CY, Menuz K & Carlson JR. (2009). Olfactory perception: receptors, cells, and circuits. Cell 139, 45–59.

31. Yuan G, Peng YJ, Khan SA, Nanduri J, Singh A, Vasavda C, Semenza GL, Kumar GK, Snyder SH & Prabhakar NR. (2016). H2S production by reactive oxygen species in the carotid body triggers hypertension in a rodent model of sleep apnea. Sci Signal 9, ra80.

32. Yuan G, Vasavda C, Peng YJ, Makarenko VV, Raghuraman G, Nanduri J, Gadalla MM, Semenza GL, Kumar GK, Snyder SH & Prabhakar NR. (2015). Protein kinase G-regulated production of H2S governs oxygen sensing. Sci Signal 8, ra37.

33. Zhou T, Chien MS, Kaleem S & Matsunami H. (2016). Single cell transcriptome analysis of mouse carotid body glomus cells. J Physiol 594, 4225–4251.

34. Zivanovic J, Kouroussis E, Kohl JB, Adhikari B, Bursac B, Schott-Roux S, Petrovic D, Miljkovic JL, Thomas-Lopez D, Jung Y, Miler M, Mitchell S, Milosevic V, Gomes JE, Benhar M, Gonzalez-Zorn B, Ivanovic-Burmazovic I, Torregrossa R, Mitchell JR, Whiteman M, Schwarz G, Snyder SH, Paul BD, Carroll KS & Filipovic MR. (2019). Selective Persulfide Detection Reveals Evolutionarily Conserved Antiaging Effects of S-Sulfhydration. Cell Metab 30, 1152–1170.e1113.

35. Zoccal DB, Bonagamba LG, Oliveira FR, Antunes-Rodrigues J & Machado BH. (2007). Increased sympathetic activity in rats submitted to chronic intermittent hypoxia. Exp Physiol 92, 79–85.

